# T cell subsets in chronic hepatitis C patients genotype 4 who achieved SVR following DAAs Therapy

**DOI:** 10.1101/2022.10.21.513286

**Authors:** Gamal Shiha, Reham Soliman, Ayman A Hassan, Nabiel NH Mikhail, Ahmed Nabil, Laila M Saleh, Doaa A Sayed, Mohamed Eslam

## Abstract

**Background:** T cells are the primary effector cells that mediate viral clearance in spontaneous recovery from HCV infection and T cell dysfunction is a hallmark of progression to chronic HCV infection.

**Material and methods:** This study included 49 well charcterised HCV genotype 4-infected patients at Egyptian Liver Research Institute and Hospital (ELRIAH), Mansoura, Egypt, who were enrolled to receive direct acting antiviral therapy for hepatitis C. Immuno-phenotyping was performed to assess the expression of multiple T cell lineage, activation and inhibitory receptors. This was done before treatment, during treatment, at end of treatment and one year after treatment. 50 patients were also enrolled as control.

**Results:** Our data showed, significant increase in the percentages of CD8+ cells as compared to control group. The percentages of PD-1 expression on the CD8+ T-cell population were signifecntly elevated in patients before treatment (p<0.001). Significant increase in Treg (CD4+CD25hFoxP3+) subsets was noticed in comparison with control pateints.

The expression of the inhibitory and activated markers in CD8+ T-cells was markedly reduced but more obvios in exhausted cytotoxic T cells compared to baseline finding (p<0.001). exhausted (PD1+CD8+) T-cells from HCV+ individuals reduced markedly after 4 weeks of DAA therapy (by 3 folds, p <0.001). Intereatingly it started to increase gradually again at the end of treatment and after 1 year but the increase doesn’t reach levels noticed in healthy control subjects.

**Conclusion:** Understanding the mechanisms of immune dysfunction and barriers to immune restoration after HCV cure will aid in better understanding of the remaining negative long-term health outcomes for HCV patients and the possibility of HCC development.

## Introduction

The World Health Organization (WHO) estimates chronic hepatitis C virus (HCV) infection prevalence to be 71 million people worldwide [1]. Chronic HCV infection (CHC) is characterized by an aberrant inflammatory response that causes HCV-mediated liver damage, leading to progressive fibrosis, potentially resulting in cirrhosis, liver failure, and hepatocellular carcinoma (HCC) [2,3].

Short-duration therapy with several oral IFN-free direct-acting antiviral agents (DAAs) has proved lead to a Sustained Virological Response (SVR) rates up to 95-100%, with minimal side effects [4]. However, achieving SVR in individuals with advanced liver disease, may only reduce but not completely eliminate the risk of developing of HCC [5–10] and a proportion of them remain on liver transplant lists [11] after HCV infection erdications.

T cells are the primary effector cells that mediate viral clearance in spontaneous recovery from HCV infection and T cell dysfunction is a hallmark of progression to chronic HCV infection [12,13]. HCV-specific CD4+ and CD8+ T cells present in CHC patients display low frequency and an exhausted phenotype characterized by a high expression of inhibitory receptors, less vigorous responses that target a limited number of epitopes of HCV [14,15], impaired cytotoxicity, reduced secretion of antiviral cytokines including IL-2 [16], IFN-λ [17,18], IL-21 [19], TNF-α [18,20], and antigen-triggered proliferation [21,22]. These dysfunctional HCV-specific T cells are characterized by a high expression of the cell surface receptor programmed cell death 1 (PD-1) exhaustion marker [23–26] that correlates with HCV viral loads [24].

Previous data during the interferon (IFN-α) era for CHC treatement suggested that IFN-α based therapy induced SVR, leads to partial, but incomplete restoration of HCV-specific immune responses. Restoration is more marked when HCV is treated during the acute phase of infection, possibly because treatment occurs before the development of exhaustion [27]. Since IFN-free DAA regimens specifically target various steps in the HCV life cycle, they can provide a unique opportunity to elucidate the interaction between HCV and the innate immune response, without the confounding effect of the IFN-α-induced immune modulation [28]. In addition, data on HCV genotype 4 is scarce.

Therefore, in this study, we aimed to evaluate the immunological changes through evaluating different T cell subsets in CHC patients who achieved SVR following DAAs therapy both at the end of treatment and after one year.

## Patients and methods

### Study Population and Ethical Considerations

This study included 49 well charcterised HCV genotype 4-infected patients at Egyptian Liver Research Institute and Hospital (ELRIAH), Mansoura, Egypt, who were enrolled to receive direct acting antiviral therapy for hepatitis C. The study protocol was approved by the local ethical committee in ELRIAH and an informed consent was obtained from each patient to be enrolled in the study. Patient characteristics and clinical parameters are provided in **Table 1.**

**Table (1):** Demographic data of 49 HCV patients.

| **Variable** | **Descriptive** |
| --- | --- |
| **Age(ys)** | 39.4±19.6 |
| **Gender** |  |
| - Females | 21 (42.8%) |
| - Males | 28 (57.2%) |
| **ALT (U/L)** | 43.07 ± 23.9 |
| **AST (U/L)** | 35.9 ± 13.4 |
| **Viral load (log<sub>10</sub> IU/ml)** | 5.57 ± 0.86 |

Quantitative variables are expressed as mean ± standard deviation
Qualitative variables were expressed as number of positive cases (%)
ALT: alanine transaminase
AST: aspartate aminotransferase

All patients were HBV surface antigen and anti-HIV negative at the time of blood collection. Patients were treated with 12 week or 24 week of daily with oral sofosbuvir (non-nucleoside polymerase inhibitor) 400 mg plus Daclatasvir (protease inhibitor) 60 mg with or without ribavirin 1000–1200 mg with dose adjustment if indicated. All treated individuals studied here achieved a sustained virological response, SVR (i.e., undetectable HCV RNA by 12 weeks after treatment cessation). Fifty apparently healthy blood donors, their ages matched with patient’s age were consider as controls.

### Labaratory investigations

Routine hematological and biochemical laboratory parameters were determined. Blood tests performed were full blood count (automated blood cell analyzer Sysmex XT-1800i (Sysmex Corporation, Kobe, Japan); liver transaminases (ALT/AST), alpha fetoprotein (AFP) (Cobas Integra 411, Roche, Basel, Switzerland), S. creatinine, S. albumin, S. bilirubin and glucose/HbA1c level (Cobas Integra 400 plus, Roche, Basel, Switzerland). HCV RNA testing was performed with real-time HCV-RNA PCR (Cobas Ampliprep, Cobas Taqman 48, Roche, Rotkreuz, Switzerland) according to manufactures instructions.

### Samples

3 ml of venous blood was withdrawn from all subjects before treatment (baseline), during treatment at week 4 (DT), at week 12 after the end of treatment at (EOT) and after one year. In addition; 2 ml blood added to EDTA tubes for flow cytometric analysis. PBMCs were isolated from whole blood by density gradient centrifugation in BD Vacutainer-CPT Mononuclear Cell Preparation tubes (BD Biosciences, San Diego, CA). PBMCs maintained in completed RPMI 1640 (GIBCO/Invitrogen) supplemented with 10% heat-inactivated human AB serum, 2 mmol/L L-glutamine and 100 μg/mL streptomycin.

### Monoclonal antibodies

Immuno-phenotyping was performed to assess the expression of multiple T cell lineage, activation and inhibitory receptors. PBMCs were stained with antibodies against surface markers, followed by staining for intracellular markers by using the following monoclonal antibodies (all purchased from Miltenyi Biotec, CA).: antihuman VioBlue conjugated anti-CD3 (REA613), FITC-conjugated anti-CD4, PE-conjugated anti-CD8, PEVio770-conjugated anti-PD-1 (CD279), APC-conjugated anti-CD38, PE-conjugated anti-CRTH-2 (CD294), PE-Vio770 conjugated anti-CXCR-3 (CD183), APC-conjugated anti-CCR-6 (CD196), PE-conjugated CD25, APC-conjugated anti-FoxP3 PE-Vio770-conjugated anti-CD127, and the corresponding isotype controls. Antibodies were used at concentrations titrated for optimal staining according to the manufacturer recommendations. Multi-colors staining of mAb were carried out as the following panels:

- CD3/CD4/CD8/CD279(PD1)/CD38 (exhausted T cells)
- CD3/ CD4/ CD294(CRTH2) /CD183(CXCR3)/CD196(CCR6) (T-helper cells)
- CD3/ CD4/ CD25/ CD127/FoxP3(T-reg)

For transcription factor detection, intranuclear permeabilization was done with eBioscience Transcription Factor Fixation/Permeabilization concentrate and diluent solutions according to the manufacturer’s instructions (eBioscience, USA).

### Flow cytometric analysis and gating strategy

Flow cytometry was performed with the MACSQuant Flow Cytometer (Miltenyi Biotec, CA). 50,000 events were acquired. Gating on viable cells then lymphocyte region (R1) based on FSC/SSC plot to exclud other cells such as RBCs, platelets aggregate and myeloid cells. Followed by gating on CD3+, subset determination based on CD4 and CD8 levels and detection of CD4/CD8 ratio. Gated T cells were sequentially gated according to the above mentioned panel for expressions of PD-1 (CD279) and CD38 to identify exhausted and activated cytotoxic T cell subsets, CXCR-3 (CD183) and CRTH-2 (CD294) to identify TH1 and TH2, CCR-6 (CD196) to identify TH17. In addition to CD25, CD127, and FoxP3 to identify Treg.

All expressions were estimated by the level of (percentage) CD4 and CD8 cell subsets from CD3 + T lymphocyts. Cut off of positivity was resolved by isotypic control.

### Statistical analysis

The data were collected, categorized and processed by using Statistical Package for Social Sciences (SPSS), version 26 software package. Data were expressed as number and percentage. Mean and standard deviation were used as a descriptive value for a quantitative data. Using paired t test was done to determine significance for numeric variables. Pearson’s correlation was also used for numeric variable.

## Results

### Changes in HCV pateints in comparison to healthy control group

#### • Increase of total CD3 T cells, CD4 and CD8 T cells in patients with HCV

Our data showed, significant increase in the percentages of CD8+ cells with no significant changes was noticed in total CD3+ and CD4+ cell type as compared to control group (Table 2). However, after treatment with DAAs the percentages of CD3+, CD4+ and CD8+ T cells were decreased to match with normal control levels (Table 3 and Fig. 1).

**Fig. 1:**
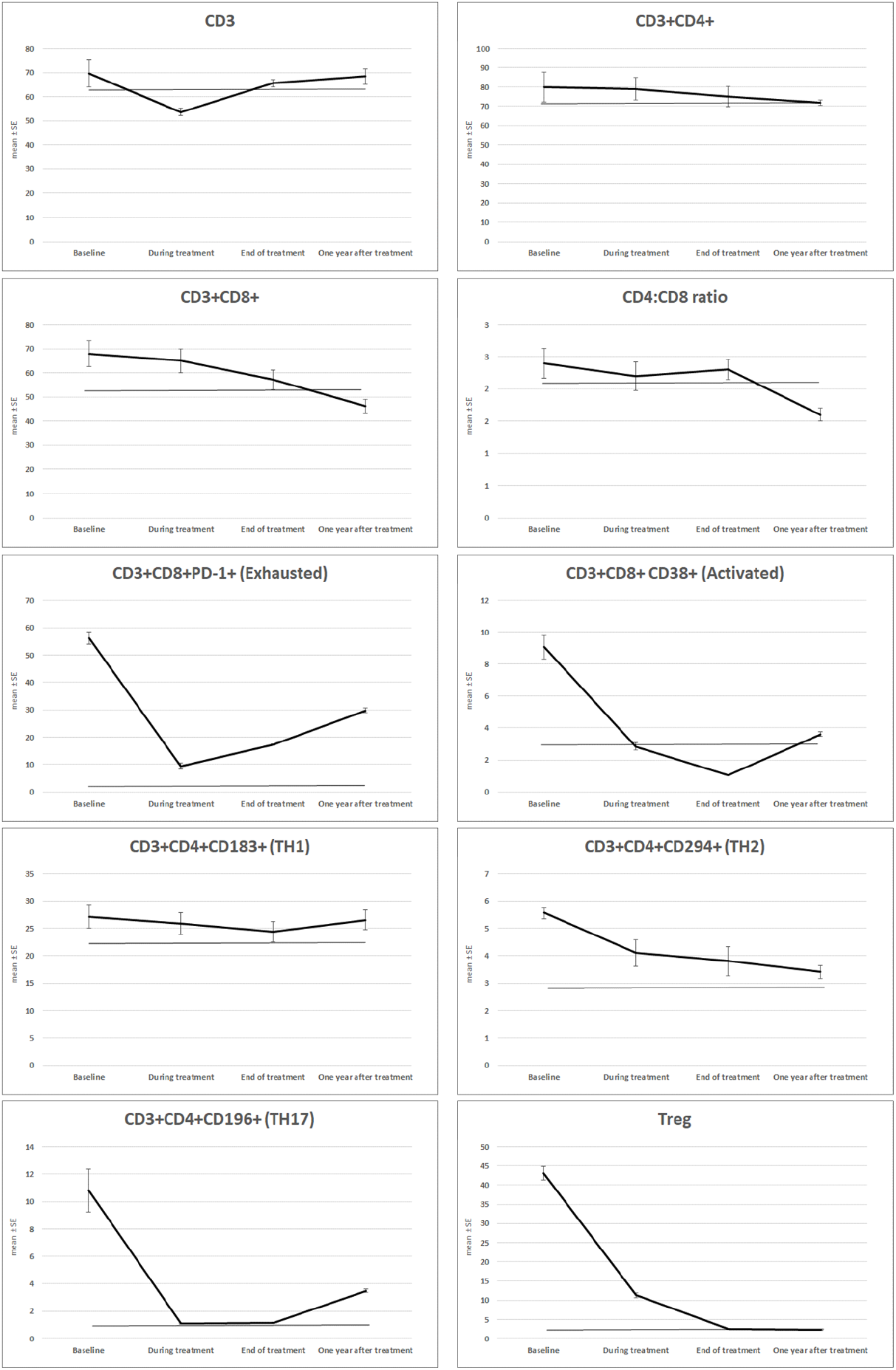
T cell subsets in chronic hepatitis C patients genotype 4 who achieved SVR following DAAs Therapy.

**Table (2):** T cell subsets in HCV patients and normal controls.

| Variables | HCV patients<br>(No.=49) | Normal control<br>(No.=50) | P value |
| --- | --- | --- | --- |
| CD3+ | 69.68 ± 39.07 | 62.22 ± 22.0 | p = 0.243 |
| CD3+CD4+ | 80.00 ± 54 | 71.0 ± 50.0 | P=0.392 |
| CD3+CD8+ | 68.0 ± 37.2 | 52.01 ± 34.00 | **P=0.028** |
| CD4:CD8 ratio | 2.4 ± 1.62 | 2.10 ± 1.5 | P=0.341 |
| CD3+CD8+ PD-1+ (Exhausted) | 56.2 ± 15.83 | 3.8 ± 1.2 | **P<0.001** |
| CD3+CD8+ CD38+ (Activated) | 9.06 ± 5.16 | 3.05 ± 0.50 | **P<0.001** |
| CD3+CD4+CD183+ (TH1) | 27.14 ± 15.1 | 22.36 ± 11.04 | P=0.075 |
| CD3+CD4+CD294+ (TH2) | 5.56 ± 1.5 | 2.8 ± 2.7 | **P=0.016** |
| CD3+CD4+CD196+ (TH17) | 10.8 ± 10.9 | 1.15 ± 0.18 | P=0.221 |
| Treg | 43.09 ± 12.75 | 2.54 ± 1.03 | **P<0.001** |

Quantitative variables are expressed as mean ± standard deviation
No: number

**Table (3):** T cell subsets in HCV patients before, during, at end of treatment and after one year.

| Variables | HCV patients<br>(baseline) | HCV patients<br>(DT) | HCV patients<br>(EOT) | HCV patients<br>after one year | P<sub>1</sub> value | P<sub>2</sub> value | P<sub>3</sub> value |
| --- | --- | --- | --- | --- | --- | --- | --- |
| CD3+ | 69.68 ± 39.07 | 53.7 ± 11.53 | 65.56 ± 10.42 | 68.34 ± 22.7 | **<0.001** | 0.8 | 0.9 |
| CD3+CD4+ | 80.00 ± 54 | 79.00 ± 41 | 75.0 ± 37.0 | 71.8 ± 10.5 | 0.14 | 0.11 | 0.07 |
| CD3+CD8+ | 68.0 ± 37.2 | 65.0 ± 34 | 57.09 ± 28.00 | 46.2 ± 20.5 | 0.44 | 0.16 | 0.09 |
| CD4:CD8 ratio | 2.4 ± 1.62 | 2.2 ± 1.54 | 2.30 ± 1.1 | 1.6 ± 0.7 | 0.14 | 0.115 | 0.08 |
| CD3+CD8+PD-1+ (Exhausted) | 56.2 ± 15.83 | 9.37 ± 7.6 | 17.26 ± 1.8 | 29.7 ± 6.2 | **<0.001** | **<0.001** | **<0.001** |
| CD3+CD8+CD38+ (Activated) | 9.06 ± 5.16 | 2.83 ± 1.64 | 1.05 ± 0.10 | 3.6 ± 1.01 | **0.001** | **<0.001** | **0.002** |
| CD3+CD4+CD183+ (TH1) | 27.14 ± 15.1 | 25.9 ± 14 | 24.36 ± 13.24 | 26.5 ± 12.6 | 0.34 | 0.32 | 0.74 |
| CD3+CD4+CD294+ (TH2) | 5.56 ± 1.5 | 4.1 ± 3.4 | 3.8 ± 3.7 | 3.4 ± 1.7 | 0.17 | 0.13 | 0.09 |
| CD3+CD4+CD196+ (TH17) | 10.8 ± 10.9 | 1.09 ± 0.4 | 1.15 ± 0.18 | 3.5 ± 0.9 | **0.001** | **0.001** | **0.002** |
| Treg | 43.09 ± 12.75 | 11.3 ± 4.1 | 2.54 ± 1.03 | 2.3 ± 1.6 | **0.002** | **<0.001** | **<0.001** |

Quantitative variables are expressed as mean ± standard deviation
P<sub>1</sub> value: p value between baseline and DT; P<sub>2</sub> value: p value between baseline and EOT; P<sub>3</sub> value: p value between baseline and 1 year after.

#### • Elevated levels of PD1+ and CD38+ CD8+ T cells in patients with HCV

We performed flow cytometric analysis of PD-1 and CD38+ on CD4+ and CD8+ T cells from HCV patients. The percentages of PD-1 expression on the CD8+ T-cell population were signifecntly elevated in patients before treatment (*p*<0.001). The increase of CD38+ on CD8+ cells was also significant (*p*<0.001) (Table 2), compared to healthy controls. The percentages of PD-1 expression on the CD8+ T-cell population showed negative correlation with age and viral load (*r*=-0.34; *p*=0.5 and r=-0.398; *p*=0.5 respectively).

#### • Increased TH17 and Treg Subsets in HCV Infection

In HCV pateints, Our flowcytometric analysis showed non significant elevations in TH17 (CD4+CD196+) and significant increase in Treg (CD4+CD25^h^FoxP3+) subsets (Table 2) in comparison with control pateints. TH1 and Treg cells showed negative correlations with age (*r*=-0.3; *p*=0.4 and *r*=-0.572; *p*<0.001 respectively), while both subsets show positive correlations with viral load (*r*=0.488; *p*=0.013 and *r*=0.584; *p*=0.002 respectively).

### Flowcytometric changes after DAAs treatment

#### • Marked decrease in exhausted and activated cytotoxic T cell subsets with treatment

To gain an insight into the kinetics by which DAA mediated HCV cure induces phenotypic changes in T cells in the peripheral blood, the detection of various T subsets was evaluated during and after treatment. The expression of the inhibitory and activated markers in CD8+ T-cells was markedly reduced but more obvios in exhausted cytotoxic T cells (Table 3 and Fig. 1) compared to baseline finding (p<0.001).

#### • Decreased of immune-regulatory T cell subsets with DAA treatment while no change in levels of TH1 and TH2 T-cell Subsets after treatment

There were significant marked reduction of the percentages of both TH17 and T reg subsets (p=0.024 and p<0.001 respectively) which had immunoregulatory functions while there were no significant differences in the the levels of TH1 and TH2 subsets of CD4 T cells during and after antiviral treatment compared to levels before treatment (Table 3 and Fig. 1).

#### • DAA therapy does not normalize PD-1+ CD8+ T-cell (exhausted cytotoxic T cells)

However, exhausted (PD1+CD8+) T-cells from HCV+ individuals reduced markedly after 4 weeks of DAA therapy (by 3 folds, p <0.001). Intereatingly it started to increase again at the end of treatment and after 1 year but the increase doesn’t reach levels noticed in healthy control subjects (Tables 3 and Fig. 1).

## Discussion

Herein, we evaluated the degree of immune restoration by performing the analysis of phenotypic changes in T cells before, during and after HCV-4 erdication by DAA therapies and in comparison to healthy control. We demonstrated significant changes in the phenotypic distribution of T-cell subsets (CD8+ subsets and T-regulatory) in the peripheral blood of CHC genotype 4 patients when compared to normal individuals.

Non significant change was noticed in CD8+ T-cells but they show decreasing trend after treatment with immediate reductions in viral load that may reflect an efflux of hepatic lymphocytes from the liver to the circulation as the virus is cleared and liver inflammation is reduced [29,30]. Pereira et al., showed that DAAs causes significant decrease of CD8+ cells post treatment in comparison to pretreatment [31]. Also, Emmanuel et al. who documented an observed decrease in CD8+ T cells in HCV patients from pretreatment to post-SVR when compared to HCV/HIV coinfected pateints suggests clearance of HCV normalizes activated T-cell levels [35].

Alanio et al. [33] found that hyperfunctional naïve CD8+ cells recovered >2 yrs post-SVR therapies. Our data (CD3+CD8+CD38+) indicates that this recovery is more early in our therapy of HCV+ individuals. So more prolonged study with clinical follow up is mandatory.

Regarding to PD-1 frequancy, our results showed significant increase of PD1+CD8+ T-cells in HCV pateints in comparison to control group. PD1+CD8+ T-cells from HCV individuals reduced markedly after 4 weeks of DAA therapy but not reach normal level. These results were in agreement with other studies that documented a significant reduction in PD-1 expression by HCV-specific CD8+ T cells following IFN-free DAAs therapy [34–36].

More interesting observation was the re-elevation of PD1+CD8+ T-cells at the end of therapy with gradual elevation after one year. In line with our results, Wieland et al. [37] mentioned that CD127+PD1+ HCV epitope-specific CD8+T cell subset is maintained during and after DAA-induced HCV clearance indicating memory like characteristcs. The CD127+PD1+ HCV-specific CD8+T cell subset is characterized by a high expression of the transcription factor-1 (TCF-1) which is a transcription factor required for the differentiation and persistence of memory CD8+ T cells. Also, CD127+PD1+ HCV-specific CD8+T cells expressed the anti-apoptotic molecule B-cell lymphoma 2 (BCL2) to agreat extent [37]. Anti-apoptotic Bcl-2 contributes to cancer formation and progression by promoting the survival of altered cells [40]. To our knomledge, it is the first time that mentioned gradual elevation of PD1+CD8+ T-cells after one year of DAAs therapy which may be a predisposing factor for hepatocellular carcinoma in chronic HCV pateints and an explanation for sustained HCC risk after viral edrication

Recently, changes in the Treg cells and Th17 cells balance were reported to be involved in disease progression and persistent hepatitis B virus (HBV) infection [38, 39]. Hence, we hypothesized the same role in HCV that an imbalance between Treg and Th17 cells participates in regulating the immune response during anti-HCV treatment. To test this possibility, we investigated the frequency of peripheral Treg cells and Th17 cells. Our results showed significant increase of Treg and Th17 in HCV pateints in comparison to control group. Pateints post DAAs treatment showed gradual dercease of Treg levels after 4 weeks of treatment then decreased to normal level at end of treatment and remains within the same level one year post treatment. [31,32]

Persistence of negative modulation by Treg cells may negatively contribute to immune reconstitution, even long after HCV has been cured, with an unknown impact on the process of carcinogenesis [41].

In conclusion, this comprehensive analysis of T cell subsets provides an expanded image of T-cell diversity at the baseline of chronic HCV infection and after response to antiviral treatment. Understanding the mechanisms of immune dysfunction and barriers to immune restoration after HCV cure will aid in better understanding of the remaining negative long-term health outcomes for HCV patients and the possibility of HCC development.

